# Molecular basis for feedback inhibition of Notch signaling by the Notch regulated ankyrin repeat protein NRARP

**DOI:** 10.1101/645937

**Authors:** Sanchez M. Jarrett, Tom C. M. Seegar, Mark Andrews, Guillaume Adelmant, Jarrod A. Marto, Jon C. Aster, Stephen C. Blacklow

## Abstract

The Notch regulated ankyrin repeat protein (NRARP) is a negative feedback regulator of Notch signaling in higher organisms. The molecular basis for NRARP function, however, has remained elusive. Mass spectrometry-based proteomic studies show that human NRARP associates with the core Notch transcriptional activation complex (NTC), containing the RBPJ transcription factor, the Notch intracellular domain (NICD), and a Mastermind-like co-activator. Binding of NRARP is direct, requires both RBPJ and NICD, and is independent of Mastermind-like proteins or DNA. The X-ray structure of an NRARP/RBPJ/NOTCH1/DNA complex, determined to 3.75 Å resolution, reveals a non-canonical mode of binding in which NRARP extends the NOTCH1 ankyrin-repeat stack by three additional repeats. Mutations of NRARP residues at the binding interface disrupt entry into complexes and suppress feedback inhibition in signaling assays. These studies establish the structural basis for NTC engagement by NRARP and provide insight into its mechanism of feedback inhibition of Notch signaling.

## Introduction

Cell signaling enables an organism to perceive and respond to its local environment. This fundamental process occurs in a series of tightly regulated steps that require stimulus detection, signal transmission, and a downstream response. The amplitude and duration of the response can be tuned by a variety of signaling modulators, which can vary widely based on the cellular context.

One common mechanism of signal modulation is feedback inhibition, in which the downstream response to the signal produces an output that suppresses the initiating signal. Feedback regulation is particularly important in developmental signaling, where control of the timing and strength of the signal is critical to ensure proper cellular proliferation and differentiation.

Notch signaling is the primary juxtacrine developmental signaling pathway controlling cell fate decisions in multicellular organisms (Bray, 2016). Mutations of the core components of this pathway give rise to a variety of developmental disorders, including Alagille syndrome (Li et al., 1997), left ventricular non-compaction (Luxan et al., 2013), spondylocostal dysostosis (Bulman et al., 2000), and Hajdu-Cheney syndrome (Isidor et al., 2011; Majewski et al., 2011; Simpson et al., 2011) as well as adult onset diseases such as CADASIL (Joutel et al., 1996). In addition, aberrant or dysregulated Notch signaling is associated with many different human cancers including T cell acute lymphocytic leukemia (T-ALL), in which activating mutations of the Notch1 receptor are found in more than half of all cases (Aster et al., 2017).

Notch signaling activation depends on cell-cell contact between a ligand-expressing “sender” cell and a receptor-expressing “receiver” cell (Fehon et al., 1990; Rebay et al., 1991). Ligand binding results in regulated intramembrane proteolysis of the receptor, liberating the intracellular portion of Notch (ICN) from the membrane (De Strooper et al., 1999; Schroeter et al., 1998; Struhl and Greenwald, 1999). ICN then migrates into the nucleus, and enters into a transcriptional activation complex that includes Notch, the transcription factor RBPJ, and a coactivator protein of the Mastermind-like (MAML) family (Petcherski and Kimble, 2000a; Petcherski and Kimble, 2000b; Wu et al., 2000), resulting in induced transcription of Notch responsive genes (see (Bray, 2016) for a review).

The Notch regulated ankyrin repeat protein (NRARP) is one of a small number of core Notch target genes. *NRARP* cDNA was first identified in an *in situ* hybridization screen following analysis of the *Delta1 synexpression* group of developmentally expressed genes in *Xenopus* embryos (Gawantka et al., 1998). The cDNA was later shown to encode a 114 amino acid protein consisting of at least two tandem C-terminal ankyrin repeats, and to be regulated at the transcriptional level by the Notch signaling pathway in *Xenopus* and mice (Krebs et al., 2001; Lamar et al., 2001).

Several studies in different organisms and developmental contexts have reported that NRARP is a negative regulator of Notch signaling. Enforced expression of *Nrarp* in mice leads to a block in T cell development, which requires Notch signaling at multiple stages (Yun and Bevan, 2003). In addition, *Nrarp* knockout mice exhibit defects in somitogenesis and vascular pruning in the eye (Krebs et al., 2012; Phng et al., 2009). Both of these phenotypes are associated with excess Notch activity, and, together with studies investigating the influence of *Nrarp* on cell fate decisions in the mouse retina (Mizeracka et al., 2013), further support the conclusion that NRARP counteracts Notch signals *in vivo*.

The molecular basis for the action of NRARP as a negative regulator of Notch signaling is incompletely understood. Experiments in *Xenopus* embryos using overexpressed tagged Notch components and NRARP detected association of NRARP with ICN and RBPJ, suggesting that the NRARP protein enters into a complex with the NTC (Lamar et al., 2001). However, proteomic studies to uncover NRARP-interacting, endogenously expressed proteins have not been performed, nor has an NRARP-NTC complex been reconstituted using purified proteins. Moreover, there are no structural data available for NRARP or NRARP-containing complexes. Thus, the molecular basis for the function of NRARP as an attenuator of Notch signal transduction has remained elusive.

Using mass spectrometry of tandem-affinity purified NRARP complexes, we show here that human NRARP associates with endogenous Notch transcription complexes (NTCs) containing Notch1, RBPJ, and a MAML coactivator. Using purified proteins, we find that NRARP binds directly to RBPJ-ICN1 complexes, requiring both RBPJ and ICN for entry into NTCs, but not MAML or DNA. The X-ray structure of a NRARP/RBPJ/ICN1/DNA complex, determined to 3.75 Å resolution, reveals that assembly of NRARP/RBPJ/Notch1 complexes relies on simultaneous engagement of RBPJ and Notch1 in a non-canonical binding mode involving the extension of the Notch1 ankyrin repeat stack by the three ankyrin repeats of NRARP. Interface mutations of NRARP disrupt entry into NTCs and abrogate feedback inhibition in Notch signaling assays. These studies establish the structural basis for NTC engagement by NRARP and provide insights into a critical negative feedback mechanism that regulates Notch signaling.

## Results

### NRARP inhibits Notch signaling and suppresses growth of Notch-dependent T-ALL cells

Previous studies in Xenopus (Lamar et al., 2001) and in mice (Krebs et al., 2012; Krebs et al., 2001; Mizeracka et al., 2013) have implicated NRARP as a negative regulator of Notch signaling (Figure 1). To test whether human NRARP inhibits Notch activity in cells, we used a well-established luciferase reporter-gene assay (Weng et al., 2003). Enforced expression of NRARP suppresses reporter activity induced by intracellular Notch1 (ICN1) in a dose-dependent manner (Figure 1B), consistent with these prior studies. To test whether the negative regulatory activity of NRARP is selective for ICN1, we also tested the effect of NRARP on reporter gene induction by ICN2, ICN3, and ICN4. The data show that NRARP inhibits reporter gene induction by all four human Notch receptors, although the inhibitory effect on ICN4- dependent reporter activity is not as strong (Supplementary Fig. S1).

**Figure 1.**
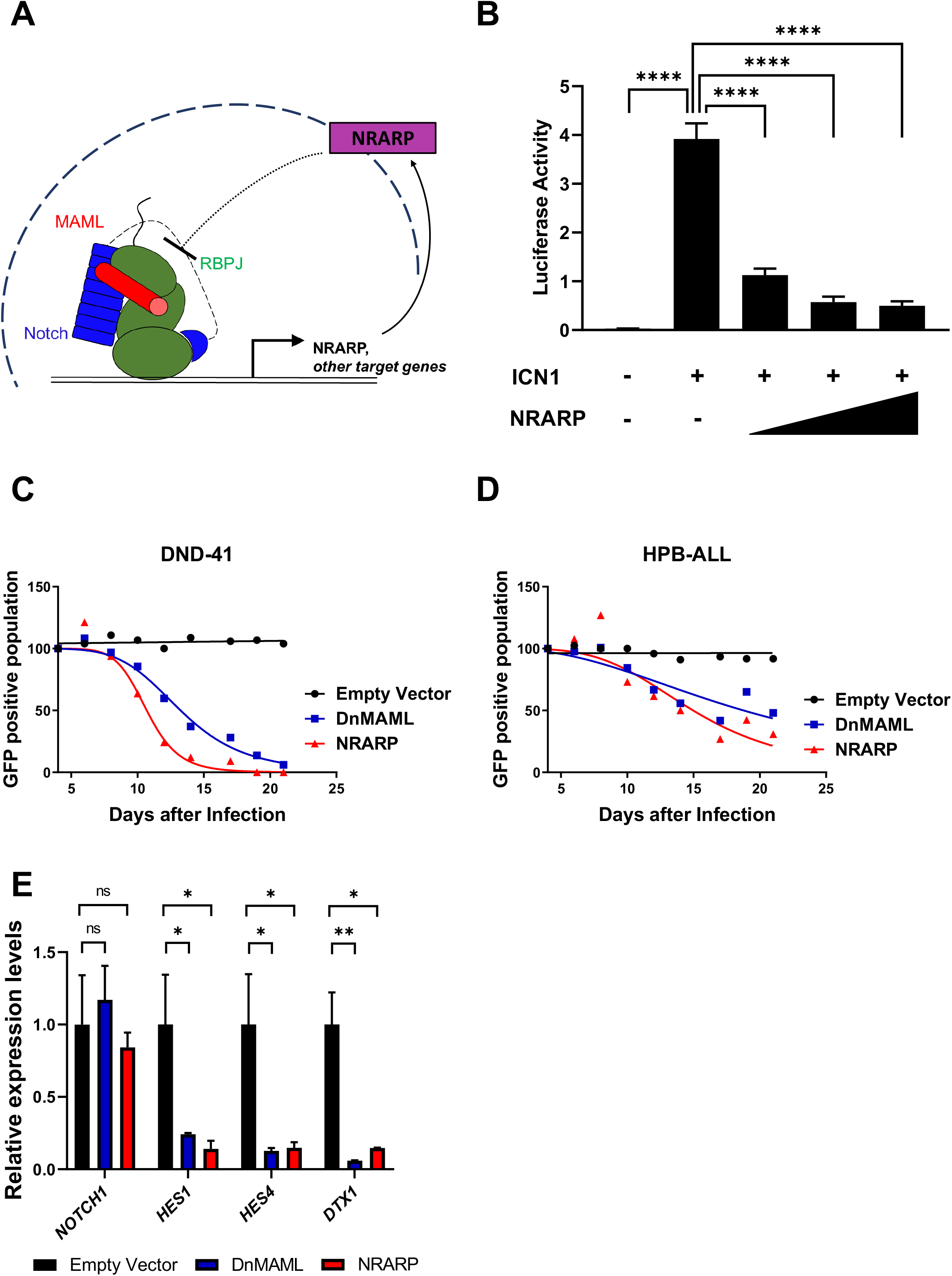
NRARP is a negative feedback regulator of Notch signaling. A. NRARP expression induced by Notch transcriptional activation complexes results in feedback repression of Notch signaling activity. B. NRARP exhibits dose-dependent inhibition of Notch signaling activity in a reporter gene assay. NIH 3T3 cells were transiently transfected with the indicated amounts of pcDNA3-based plasmids for expression of ICN1 and NRARP, a firefly luciferase reporter plasmid under control of the TP1 Notch response element, and a plasmid expressing Renilla luciferase. Firefly luciferase activity is reported relative to Renilla control, setting this ratio for cells transfected with empty pcDNA vector to a value of one. Statistical analysis was performed using ANOVA, and a Dunnett’s multiple comparison post hoc test was performed comparing test samples to the control. ****p < 0.0001. C, D. Effect of NRARP or dnMAML on the growth of DND-41 (C) and HPB-ALL (D) T-ALL cells. Each cell line was transduced with retrovirus expressing dnMAML1-GFP, NRARP with GFP behind an IRES, or GFP alone. The relative fraction of GFP positive cells is plotted as a function of time. E. RNA abundance of *NOTCH1* and sentinel Notch target genes *HES1*, *HES4*, and *DTX* in NRARP transduced Jurkat cells, measured using qPCR. Statistical analysis was performed using ANOVA, and a Dunnett’s multiple comparison post hoc test was performed comparing test samples to the control. *p < 0.05; **p < 0.01.

Previous studies have shown that genetic and chemical inhibition of Notch in Notch-mutated T-ALL cell lines results in suppression of cell growth. We tested whether enforced expression of NRARP in two Notch-dependent T-ALL cell lines, DND-41 and HPB-ALL, results in growth suppression, as predicted for a negative modulator of signaling. Cells were infected with GFP-expressing retroviruses that were either empty or carried the cDNA for *NRARP* or dominant-negative *MAML1* (Weng et al., 2003), and the GFP positive populations of unsorted transduced cells were monitored over time to assess the effect of viral transduction on cell growth. Transduction with *NRARP* results in suppression of growth of both T-ALL lines to a similar extent as dominant-negative *MAML1* (dnMAML), whereas empty virus has no effect (Figure 1C, D). Analysis of the Notch-responsive genes *HES1*, *HES4*, and *DTX1* in cells expressing *NRARP* revealed decreased levels of mRNA (Figure 1E), again consistent with the conclusion that NRARP is a negative regulator of Notch signaling. The amount of *NOTCH1* mRNA, on the other hand, is unchanged in *NRARP*-expressing cells, suggesting that NRARP regulates NOTCH1 not by influencing its expression, but by modulating its activity at the protein level (Figure 1E).

### Direct binding of NRARP to NOTCH1-RBPJ complexes requires both RBPJ and NOTCH1

Previous work carried out in *Xenopus* embryos using forced expression of tagged proteins showed that NRARP co-immunoprecipitates with Xenopus RBPJ and Xenopus ICN1, suggesting that NRARP enters a complex with Notch and RBPJ (Lamar et al., 2001). To identify the complete spectrum of proteins that associate with human NRARP, we used tandem affinity purification of HA-FLAG-tagged NRARP from Jurkat cells followed by mass spectrometry of the recovered endogenous proteins. We consistently recovered peptides for the three core components of the NTC: RBPJ, NOTCH1, and MAML1, in four independent experiments (Table 1). A complete list of all proteins detected in each of these experiments is presented in Supplementary Table 1.

**Table 1.**
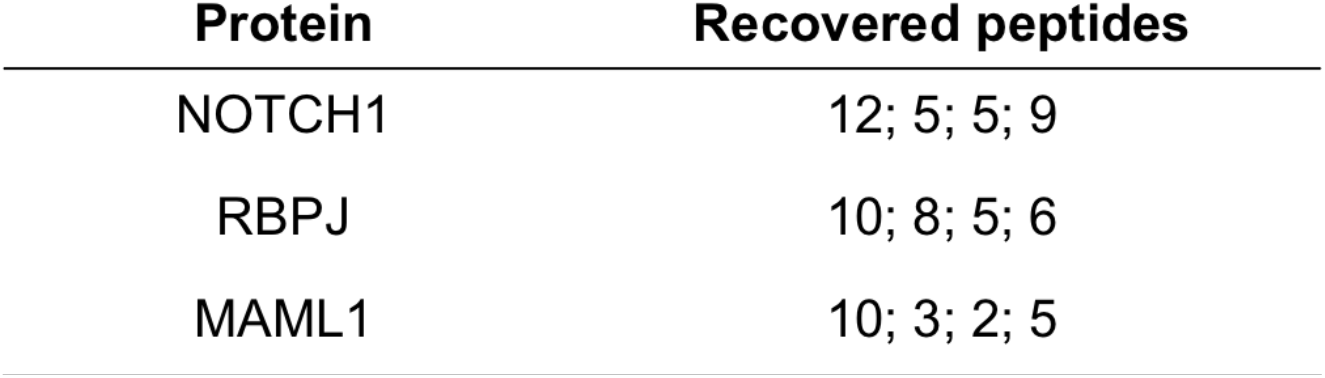
Tandem immunoprecipitation and mass spectrometry results from Jurkat cells using tagged NRARP as bait. The number of unique endogenous peptides recovered for components of the core Notch transcription activation complex is listed for 4 independent experiments.

RBPJ contains three structured domains that encompass most of the coding sequence, followed by a region that is not required for assembly of RBPJ-ICN1-MAML1 complexes on DNA. ICN1 has a RAM region, a series of ankyrin repeats (ANK), a transcriptional activation region (TAD), and a C-terminal PEST sequence. NRARP is predicted to have three ankyrin repeats, and MAML1 is predicted to be unstructured C-terminal to the region required for formation of the NTC (Figure 2A).

**Figure 2.**
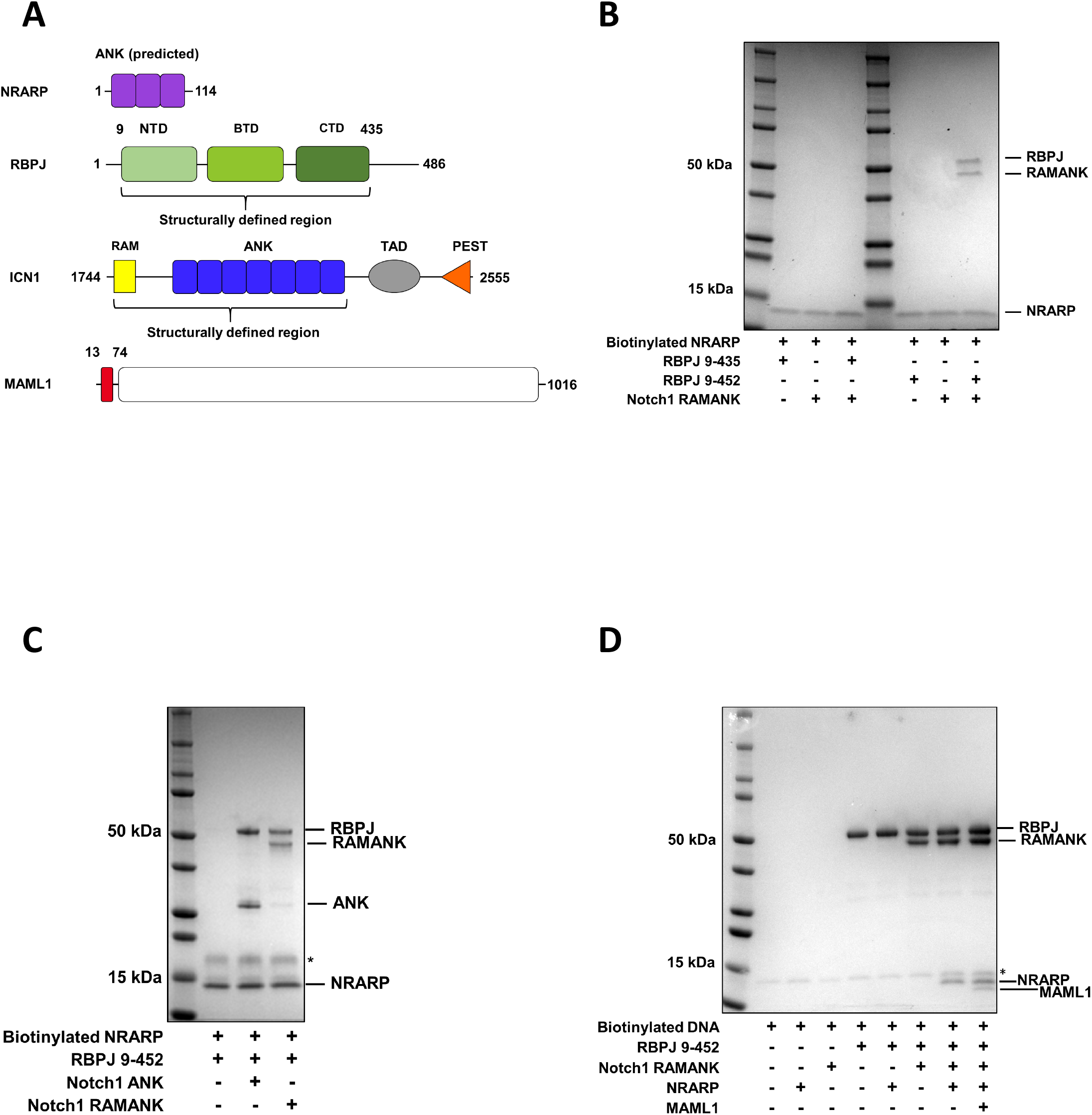
Requirements for complexation of NRARP with Notch1 and RBPJ. A. Domain organization of the protein components found in NRARP complexes. NRARP has three predicted ankyrin-repeats (ANK). RBPJ contains an N-terminal domain (NTD), β-trefoil domain (BTD), and a C-terminal domain (CTD). Intracellular Notch1 (ICN1) contains an RBPJ associated molecule (RAM), an ankyrin-repeat domain (ANK), a transcriptional activation domain (TAD), and a PEST (proline, glutamate, serine and threonine) sequence. MAML1 contains an N-terminal Notch/RBPJ binding region (red) followed by a C-terminal portion predicted to be natively disordered (white). B. Binding of NRARP requires both RBPJ and Notch1, and a C-terminal RBPJ extension. Biotinylated-NRARP, RAMANK, and the indicated forms of RBPJ were combined, complexes were recovered using streptavidin-sepharose beads, and the recovered proteins were analyzed on a Coomassie-stained gel. C. The ANK domain of Notch1 is sufficient for recruitment of NRARP into RBPJ-Notch1 complexes. Biotinylated-NRARP, RBPJ and either ANK or RAMANK were combined, complexes were recovered using avidin-sepharose beads, and the recovered proteins were analyzed on a Coomassie-stained gel. D. Association of NRARP with Notch1/RBPJ complexes is compatible with both MAML1 and DNA binding. Biotinylated-DNA (containing a single RBPJ binding site), MAML1(13-74), NRARP, RBPJ and Notch1 RAMANK proteins were combined, complexes were recovered using streptavidin-sepharose beads, and the recovered proteins were analyzed on a Coomassie-stained gel.

To map the requirements for formation of complexes between NTC components and NRARP, we purified the RAM-ANK region of NOTCH1 and the structured portion of RBPJ. Neither of these proteins, alone or in combination, form stable complexes with NRARP (Figure 2B). Remarkably, however, extension of the C-terminus of RBPJ to residue 452 enabled purification of stable complexes, but only when both RBPJ and Notch1 are present (Figure 2B). Further domain-mapping studies established that the only region of ICN1 that is required for complex formation is the ANK domain (Figure 2C), and that association of NRARP with NOTCH1 and RBPJ does not compete with either MAML1 or DNA binding (Figure 2D),

### Crystal structure of an NRARP-NOTCH1-RBPJ-DNA complex reveals a composite binding interface

To determine the structural basis for recognition of NOTCH1-RBPJ complexes by NRARP, we determined a 3.75 Å X-ray structure of an NRARP-NOTCH1-RBPJ complex bound to DNA, phased using molecular replacement with the human NOTCH1/RBPJ/MAML1/DNA complex (PDB ID code 2F8X; (Nam et al., 2006)) as a search model (Table 2). The structural features described here are drawn from the better ordered of the two assemblies seen in the asymmetric unit (Supplementary Figure 2A, B).

**Table 2.** Data collection, structure determination and refinement statistics.

| Data Collection | NRARP-NTC |
| --- | --- |
| Wavelength ($\lambda$, Å) | 0.979 |
| Resolution range (Å) | 48.46 - 3.754 (3.888 - 3.754) |
| Space group | P2<sub>1</sub>2<sub>1</sub>2<sub>1</sub> |
| Unit cell (Å, degrees) | 79.88, 103.65, 301.42, 90, 90, 90 |
| Total reflections | 114,867 (9,192) |
| Unique reflections | 26,271 (2,504) |
| Multiplicity | 4.4 (3.7) |
| Completeness (%) | 99.17 (97.13) |
| Mean I/sigma(I) | 7.39 (1.16) |
| Wilson B-factor | 114.54 |
| R-merge | 0.174 (1.155) |
| R-meas | 0.1973 (1.343) |
| CC<sub>1/2</sub> | 0.998 (0.689) |
| Reflections used in refinement | 26,221 (2,502) |
| Reflections used for R-free | 1,997 (190) |
| R-work | 0.267 (0.360) |
| R-free | 0.315 (0.402) |
| Number of non-hydrogen atoms | 13,310 |
| Macromolecules | 13,310 |
| Protein residues | 1,551 |
| RMS (bonds, Å) | 0.002 |
| RMS (angles, degrees) | 0.58 |
| Ramachandran favored (%) | 92.4 |
| Ramachandran allowed (%) | 7.3 |
| Ramachandran outliers (%) | 0.3 |
| Rotamer outliers (%) | 0 |
| Clashscore | 10.1 |
| Average B-factor (Å<sup>2</sup>) | 119.4 |

\*Highest shell statistics are reported in parentheses.

The most striking feature of the complex is the assembly of the ankyrin repeats from NOTCH1 and NRARP into a pseudo-continuous stack that wraps around the RBPJ-DNA complex in a crescent-shaped arc (Figure 3A). The extended ankyrin repeat stack results in an elongated assembly overall, with dimensions of approximately 120 × 70 × 60 Å. The arrangement of RBPJ, NOTCH1, and the DNA within the complex are minimally affected by the binding of NRARP, as the NOTCH1 and RBPJ subunits of the NRARP complex superimpose with a backbone root-mean-square deviation (RMSD) of 1.06 Å when compared with the transcriptional activation complex that contains NOTCH1, RBPJ and MAML on DNA (Figure 3B, C).

**Figure 3.**
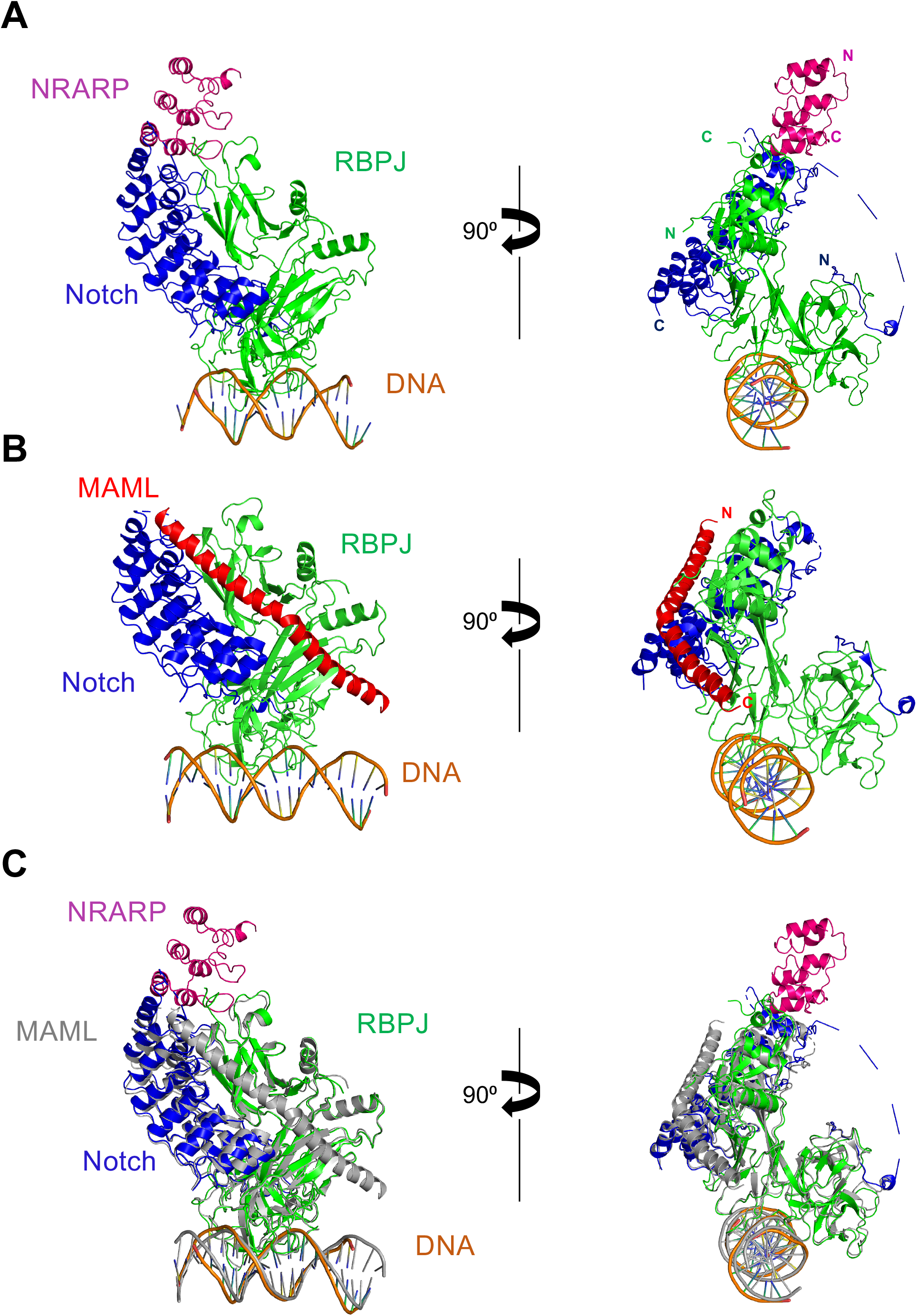
Structure of an NRARP-Notch1-RBPJ complex on DNA, and comparison with the human Notch1-RBPJ-MAML1 (PDB ID code 3V79; (Choi et al., 2012)) transcriptional activation complex. A. Ribbon representation of the NRARP-Notch1-RBPJ complex on DNA. The complex contains NRARP (pink), RBPJ (green), Notch1 RAMANK (blue) and a 16-mer DNA (orange) containing a single RBPJ binding site. B. X-ray structure of RBPJ in complex with the ankyrin domain of Notch1 (blue) and MAML1 (red) on DNA (orange). C. Overlay of the NRARP-Notch1-RBPJ-DNA complex (colors) with the RBPJ-Notch1-MAML1-DNA complex (gray).

NRARP itself is a single protein domain with three ankyrin repeats. Even in the structure of the complex, however, the first ankyrin repeat is less ordered that the other two, and its first helix is modeled only as polyalanine even in the better-defined copy of the asymmetric unit. The third ankyrin repeat of NRARP engages the NOTCH1-RBPJ interface at a composite surface that includes the first ankyrin repeat of NOTCH1 and the C-terminal domain of RBPJ. At this interface, NRARP is oriented with its C-terminal ankyrin repeat abutting the N-terminal ankyrin repeat of NOTCH1, thereby creating the pseudo-continuous stack of repeats that travels along the C-terminal Rel-homology domain of RBPJ (Figure 3A). The NRARP-NOTCH1 interface results in improved electron density for the first ankyrin repeat of NOTCH1 when compared with the structure of the human NTC, suggesting that NRARP binding stabilizes the structure of this repeat. The interaction between NRARP and RBPJ relies on the canonical concave binding surface of the third ankyrin repeat of NRARP, which appears to approach within contact distance of the C-terminal extension of RBPJ. The interface between NRARP and the NOTCH1-RBPJ complex is non-overlapping with the NOTCH1-RBPJ interface with MAML1, and completely compatible with the observed simultaneous binding of NRARP and MAML1 by RBPJ-Notch complexes on DNA (Figure 3C). Moreover, the NRARP binding site is also non-overlapping with the NTC dimerization interface (Arnett et al., 2010) on the convex face of the NOTCH1 ankyrin domain (Supplementary Figure 2C). Key NRARP residues at the contact interface with NOTCH1-RBPJ include W85 and A92 of the third ankyrin repeat (Figure 4A). W85 makes contacts in a cleft created primarily by residues on RBPJ, with an additional potential contact with P1880 of NOTCH1, whereas A92 (stick) of NRARP approaches the first helix of the NOTCH1 ANK domain (Figure 4B).

**Figure 4.**
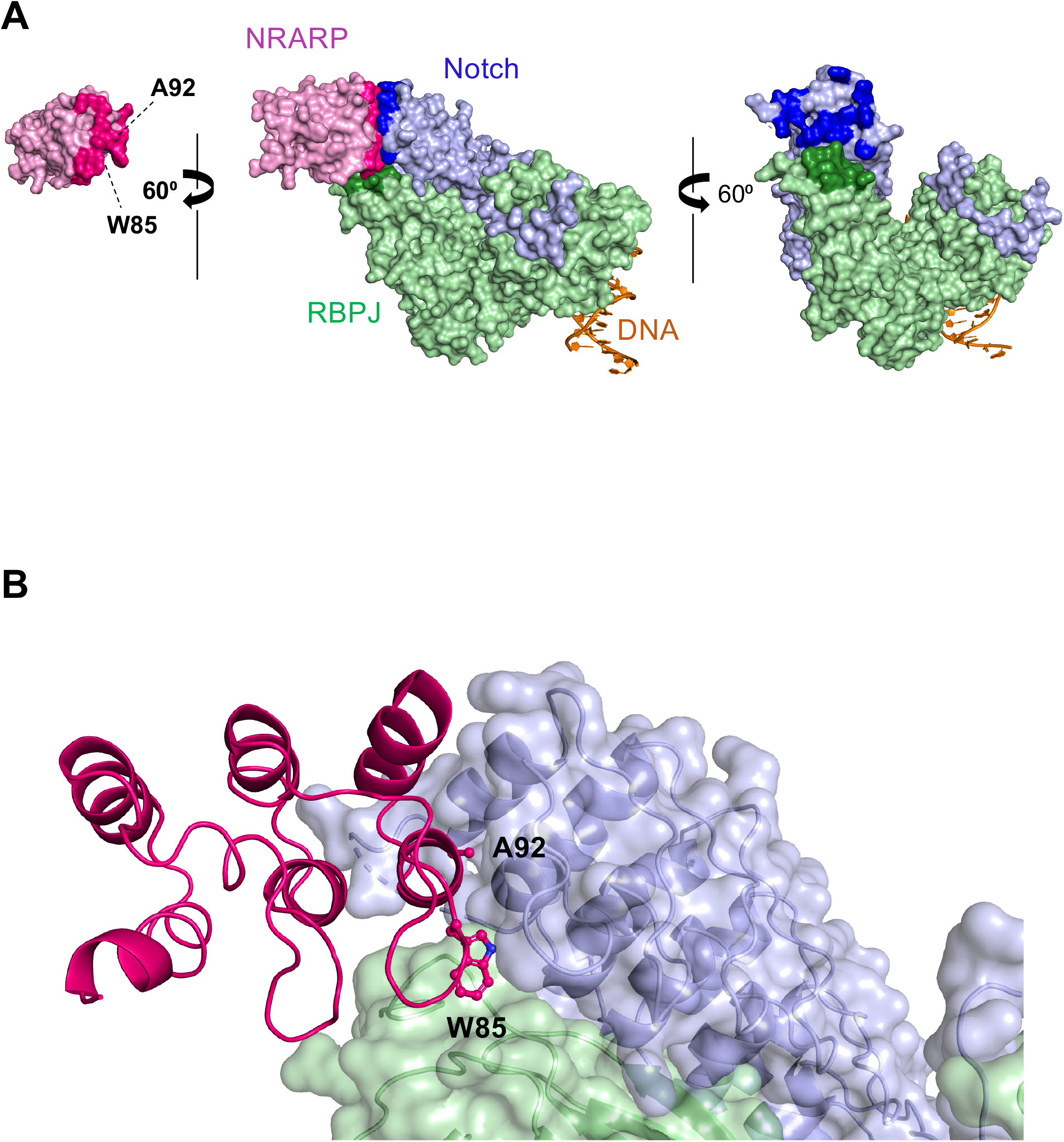
**A.** Molecular surface representation highlighting interactions between NRARP and the Notch1-RBPJ interface. Center panel: NRARP is pink, RBPJ is green and the ANK domain of Notch1 is blue. Left panel: open book view of NRARP. NRARP is rotated 60 degrees clockwise and residues that approach within 4 Å of the ANK-RBPJ surface are colored in a darker shade. Interface residues W85 and A92 are indicated. Right panel: open book view of the ANK-RBPJ surface. ANK-RBPJ is rotated 60 degrees counterclockwise and residues that approach within 4 Å of NRARP are colored in darker shades. B. Close-up view of the binding interface. W85 of and A92 of NRARP are shown in ball and stick form.

### Inhibition of Notch signaling by NRARP depends on the binding interface

To determine whether the inhibition of Notch activity by NRARP relies on the binding interface seen in the X-ray structure, we tested the effect of mutating conserved NRARP residues at this interface in the reporter gene assay. The first mutation, W85E, significantly attenuates the inhibitory effect seen with wild-type NRARP. When combined with an additional A92W mutation (W85E/A92W), the attenuation is even greater (Figure 5A). Neither the single nor the double mutation disrupt the overall structural integrity of purified NRARP protein (Figure 5B), as judged by the near equivalence of their circular dichroism (CD) spectra (Figure 5C). To determine whether the reduced inhibitory activity of the NRARP mutants is indeed due to a decrease in NRARP binding as predicted, we directly tested binding of purified recombinant NRARP polypeptides to RBPJ-NOTCH1 complexes on DNA. Whereas wild-type NRARP enters NOTCH1-RBPJ complexes on DNA, neither W85E nor W85E/A92W do so, confirming that both mutants are defective in forming complexes (Figure 5D).

**Figure 5.**
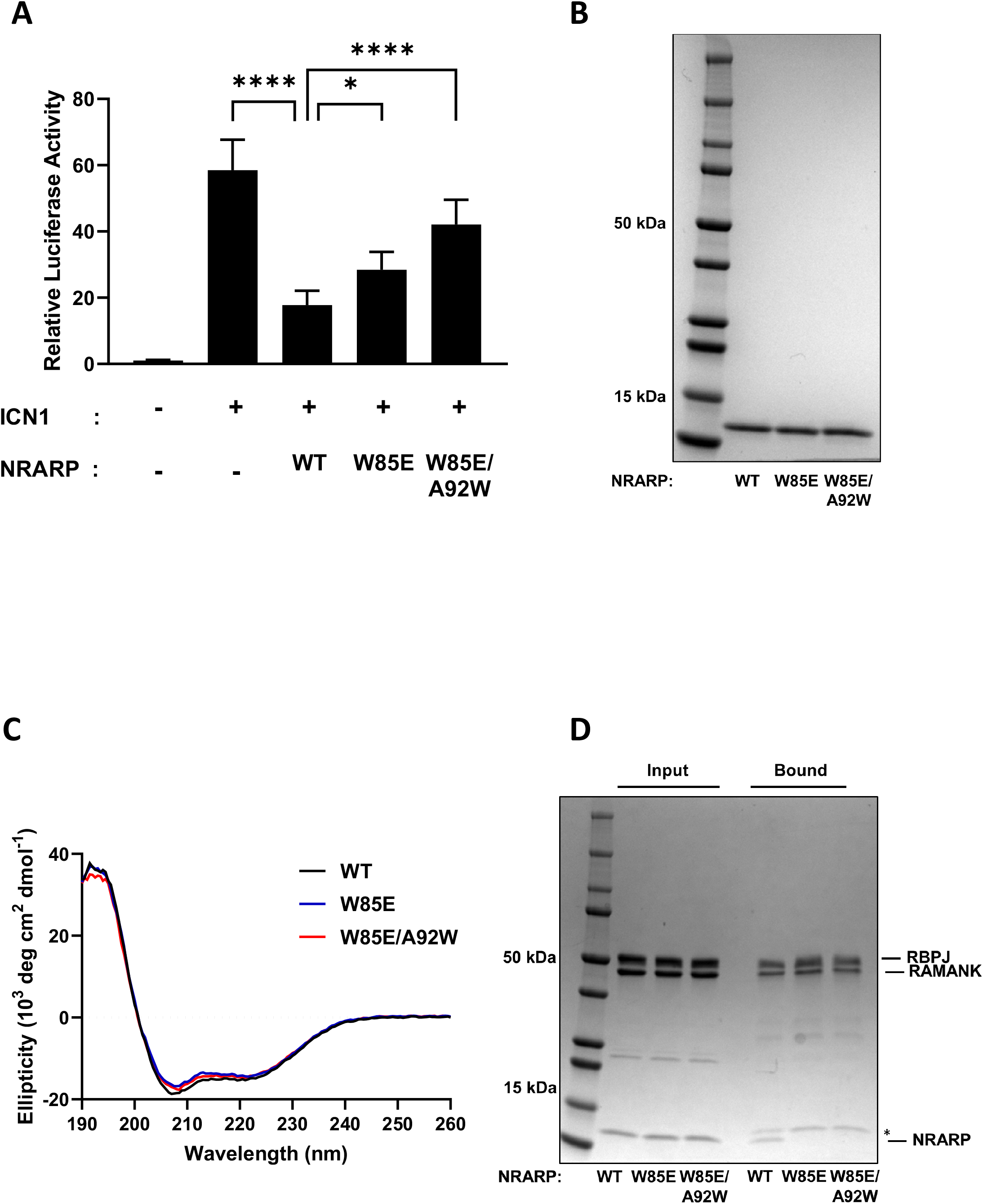
A. Effect of NRARP interface mutations on Notch-dependent luciferase reporter gene activity. Statistical analysis was performed using ANOVA, and a Dunnett’s multiple comparison post hoc test was performed comparing test samples to the control. *p < 0.05; ****p < 0.0001. B. Coomassie-stained gel of purified wild-type (WT), W85E, and W85E/A92W forms of NRARP. C. CD spectra of purified WT, W85E, and W85E/A92W NRARP proteins. D. Effect of NRARP interface mutations on binding to preassembled Notch1-RBPJ complexes, captured on biotinylated DNA.

### NRARP promotes NOTCH turnover

Prior studies have reported a decrease in detectable levels of ICN when both NOTCH1 and NRARP are transiently co-expressed in *Xenopus* embryonic extracts (Lamar et al., 2001). To determine whether NRARP affects the abundance of endogenous Notch1 in human cells, we infected Jurkat cells with control retrovirus expressing GFP only, virus expressing dnMAML1, or virus expressing NRARP, and probed cell lysates for both total Notch1 and activated Notch1 (ICN1). Whereas expression of dnMAML leads to accumulation of activated Notch1 (Figure 6A) compared to vector control, expression of NRARP reduces the level of ICN1 without depleting total Notch1 (Figure 6A), suggesting that NRARP selectively promotes degradation of the active intracellular form of Notch1 (ICN1).

**Figure 6.**
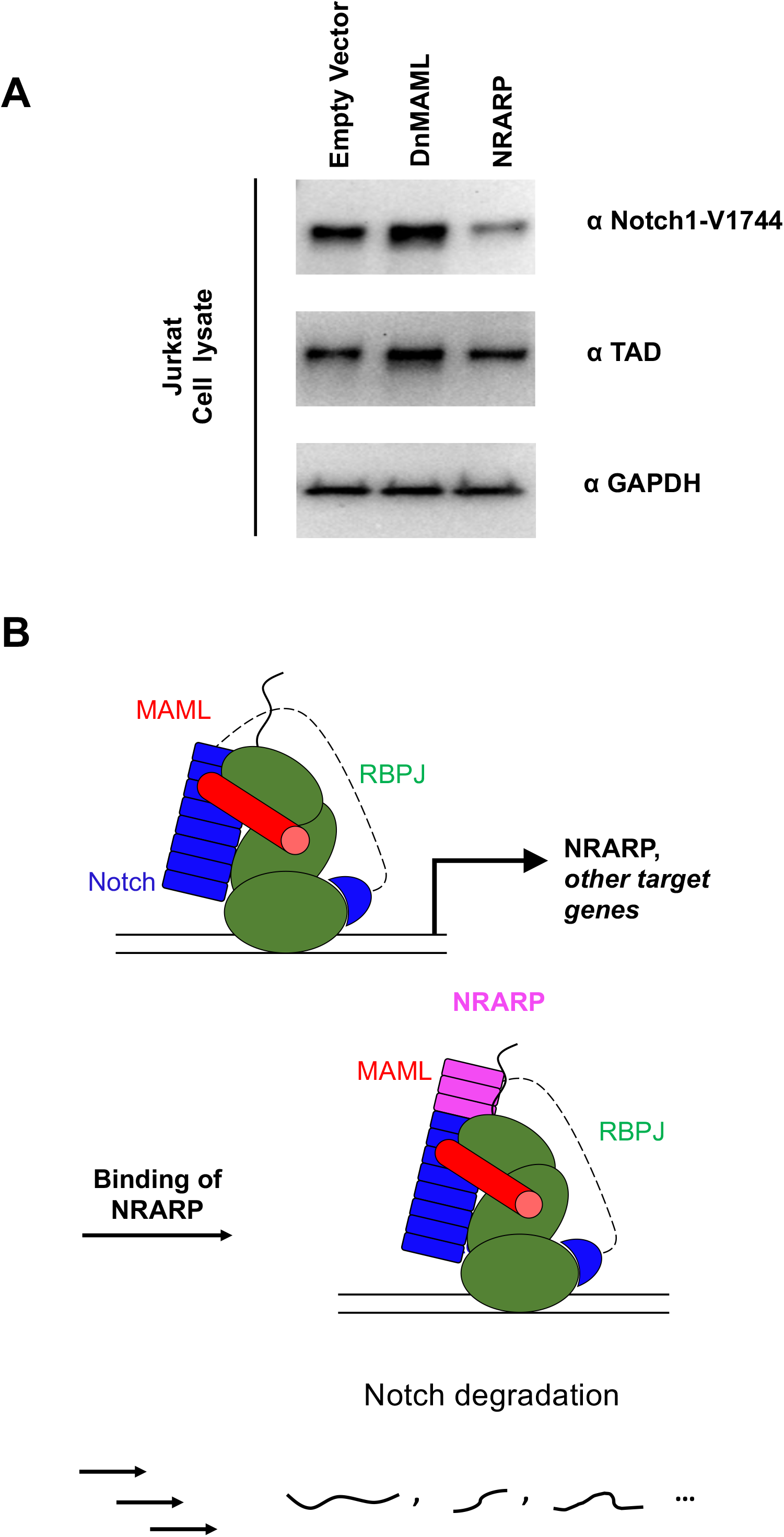
NRARP promotes degradation of activated Notch1 complexes. A. Jurkat cells were transduced with retrovirus expressing GFP alone, a GFP-DnMAML fusion protein, or NRARP, and the amounts of activated Notch1 (top panel), total Notch1 (middle panel), or GAPDH (bottom panel) were determined by Western blot with the indicated antibodies. B. Proposed model for NRARP function in Notch signaling. Notch pathway activation results in transcription of NRARP, a Notch target gene. NRARP binds to Notch transcriptional activation complexes, accelerating degradation of intracellular Notch and downregulating target gene expression.

## Discussion

The induced expression of Notch target genes relies on the formation of an NTC containing the transcription factor RBPJ, ICN, and a MAML coactivator on DNA. The stepwise assembly of an NTC begins with ICN binding to RBPJ, which enables the engagement of MAML followed by the recruitment of additional co-factors for target gene transcription. RBPJ also mediates transcriptional repression in the absence of Notch by interacting with an alternative set of cofactors, including several co-repressors, such as SHARP (Kuroda et al., 2003; Oswald et al., 2002) and KyoT2 (Taniguchi et al., 1998).

We report here the structure of an NRARP-NOTCH1-RBPJ complex bound to DNA, revealing the molecular basis for interaction of the Notch feedback inhibitor NRARP with the core NTC. Previous work anticipated the direct interaction of NRARP with Notch-RBPJ complexes from studies with overexpressed proteins in *Xenopus* extracts (Lamar et al., 2001). Consistent with those observations, we detected a robust interaction between human NRARP and endogenous components of the core human NTC in Jurkat cells using an unbiased proteomic approach. When we reconstituted a complex with purified proteins, we found that NRARP associates directly with Notch-RBPJ complexes and requires the presence of both Notch and RBPJ for complex formation, much as MAML proteins require both Notch and RBPJ for NTC assembly. Our work with purified proteins also established that a C-terminal extension of RBPJ beyond the region used in previous structural studies is required for stable entry of NRARP into RBPJ-Notch complexes, but that neither MAML nor DNA is needed for NRARP recruitment.

NRARP binding does not induce any major conformational changes in the Notch-RBPJ complex, nor does it interfere with binding of MAML1 or DNA. Instead, NRARP extends the ankyrin repeat stack of Notch1 by three repeats, engaging the first of the NOTCH1 ankyrin repeats with its C-terminal one. NOTCH4, which is the human homolog least sensitive to inhibition in reporter gene assays (Supplemental Figure 1), also shows the greatest divergence in its first ankyrin repeat, explaining its reduced sensitivity to NRARP inhibition. In contrast, NRARP relies on the concave surface of its ankyrin repeat stack to contact RBPJ, using a binding mode seen frequently in other ankyrin repeat protein complexes. The induced ordering of the C-terminal extension of RBPJ forces the serine/threonine-rich C-terminal segment of RBPJ away from the core of the NTC, potentially exposing it to post-translational modifications that may regulate NTC turnover.

Because NRARP binds to a composite RBPJ-Notch surface, its action is restricted to effector signaling complexes that are engaged in inducing a transcriptional response. Prior work (Fryer et al., 2004) pointed to a link between the assembly of Notch transcription complexes to their timed destruction, with estimates for the half-life of activated (gamma-secretase-cleaved) Notch of roughly 2-4 h. Transcriptional induction of canonical target genes like NRARP in response to activated Notch can occur within 1 h, indicating that negative feedback regulation by direct binding of NRARP to promote Notch turnover may be one of the early steps in the molecular mechanism underlying this “timed destruction” program.

A number of mechanistic explanations may account for why binding of NRARP to the NTC inhibits Notch target gene activation. One possibility is that NRARP binding directly alters the ability of the NTC to recruit co-factors to stimulate transcription, but this explanation seems unlikely because NRARP does not appreciably affect either DNA or MAML binding. Another possibility is that bound NRARP recruits enzymes that directly modify the NTC to suppress transcriptional induction. Along these lines, there are phosphorylation sites near the NRARP binding site on NOTCH1 (Ranganathan et al., 2011). Though these sites (T1898 and S1901) are distant from the DNA contact interface on RBPJ and unlikely to directly affect DNA binding when modified, their phosphorylation could indirectly affect the ability of NTCs to stimulate transcription. The most likely model for NRARP action, however, supported both by data presented here (Figure 6A) and by complementary studies using *Xenopus* extracts (Lamar et al., 2001), is that NRARP accelerates NTC turnover (Figure 6B), likely by promoting such posttranslational modifications of Notch, as well as of RBPJ and/or MAML. The molecular pathway for NRARP-mediated NTC turnover, which appears to be present in both physiological and pathological contexts, could be a future avenue for development of therapeutics designed to modulate the dynamic response of cells to Notch pathway activation.

## Materials and Methods

### Protein expression and purification

Wild type and mutant full length NRARP constructs were assembled into a pETHSUL vector at BamHI and XhoI sites to produce His-SUMO fusion molecules. A cassette encoding full-length biotinylated NRARP was assembled by placing a biotinylation (avi) tag between the SUMO tag and NRARP. This cassette was inserted into a pETDUET vector encoding biotin ligase (BirA) for NRARP expression and in-cell biotinylation. Point mutants were constructed by site-directed mutagenesis.

For protein production, expression constructs were transformed into Rosetta(DE3)pLysS cells and cells were grown in rich media at 18°C. Protein expression was induced with 0.5 mM isopropyl β-D-1 thiogalactopyranoside (IPTG), and cells were grown for an additional 16 h at 18°C. Cells were harvested by centrifugation, subjected to a freeze-thaw cycle, and lysed by sonication in 50 mM Tris buffer, pH 8.0, containing 500 mM NaCl, 5% glycerol, and 2mM tris(2-carboxyethyl)phosphine (TCEP), and supplemented with EDTA-free protease inhibitor tablets (Roche).

NRARP protein was affinity-captured from cleared lysates by affinity chromatography using HisPur Ni-NTA resin (Thermo Scientific). NRARP was cleaved from the His-SUMO tag and released from the beads using Ulp1 protease. The NRARP protein was then purified by anion exchange chromatography on Mono-Q resin using a linear gradient of NaCl (0.05-1 M) in 20 mM Tris buffer, pH 8.0, containing 1 mM EDTA and 5 mM DTT. Fractions containing the partially purified NRARP protein were combined, concentrated and further purified by size-exclusion chromatography using a Superdex 200 column equilibrated in 20 mM Tris buffer, pH 8.5, containing 500 mM NaCl, 5% glycerol and 2 mM TCEP. Fractions that were >95% pure as assessed by SDS-PAGE were pooled, flash frozen, and stored at −80° C.

RBPJ molecules (9-452 and 9-435) were expressed and purified using the same series of chromatographic steps. The only modification was the buffer used for anion exchange on Mono-Q resin. Because RBPJ is not stable in low-salt buffer, the linear gradient of NaCl was from 0.1-1 M in 20 mM Tris buffer, pH 8.3, containing 5 mM DTT. MAML1, Notch1 ANK and Notch1 RAMANK proteins were expressed and purified as previously reported (Nam JBC).

### Crystallization and Data collection

The components used to generate protein complexes for crystallography were full-length human NRARP, residues 1760-2126 of human Notch1 (comprising the RAM and ANK domains) and residues 9-452 of human RBPJ. Complexes were purified by size exclusion chromatography using a Superdex 200 column equilibrated in 10 mM Hepes buffer, pH 7.8, containing 150 mM NaCl and 2 mM TCEP. A 16mer oligomeric DNA duplex containing an RBPJ binding site with two nucleotide overhangs was generated by annealing oligonucleotides (5’-TTGACTGTGGGAAAGA-3’, 5’-AATCTTTCCCACAGTC-3’) at 95 °C for 5 minutes and slowly cooling to room temperature. The trimeric protein complex and DNA were combined in a ratio of 1:1.1. Crystals of protein-DNA complex (4 mg/ml) grew in sitting drops at 16 °C after 24-36 hours in 50 mM Hepes pH 6.8, 200 mM Sodium Fluoride, 18% PEG3350, Crystals were cryoprotected by supplementing the mother liquor with 20% ethylene glycol (v/v). Data were collected at the Advanced Photon Source, beamline 23-ID-B (GM/CA).

### Structure Determination

The structure was solved by molecular replacement in Phenix with Phaser (McCoy et al., 2007), using RBPJ and the NOTCH1 ANK-domain from the human NTC structure, PDB 2F8X (Nam et al., 2006). The presence of density for DNA and for the RAM portion of NOTCH1 confirmed that the molecular replacement solution was correct. Two NRARP/NOTCH1/RBPJ/DNA complexes were identified in the asymmetric unit. Iterative rounds of manual model building and refinement were performed with COOT (Emsley and Cowtan, 2004) and phenix.refine (Afonine et al., 2012), respectively. All crystallographic data processing, refinement, and analysis software was compiled and supported by the SBGrid Consortium (Morin et al., 2013). PyMOL (Schrodinger) was used to prepare all figures as well as perform structural superpositions. Coordinates have been deposited in the protein data bank with PDB ID code xxx.

### Reporter assay

NIH 3T3 cells cultured in 24-well plate format were transfected with pcDNA3-based plasmids for 100 ng of ICN1, a varying amount of NRARP (as indicated in Figures 1(0.5-2 μg) and 5 (500ng)), a firefly luciferase reporter plasmid under control of the TP1 Notch response element, and a Renilla luciferase control plasmid. Cells were harvested 24 hours after transfection. The firefly luciferase activity, relative to Renilla control was measured using the Dual Luciferase kit (Promega). Each measurement is plotted normalized to the signal from 3T3 cells transfected with empty pcDNA3 vector, which was set to a value of 1.

### Cell growth assay

Each cell line was transduced with retrovirus encoding NRARP with expression linked to GFP via an IRES, MAML1(13-74)-GFP fusion or GFP retrovirus at titers such that only a subpopulation of cells was transduced. The cell population was identified daily by flow cytometry using forward/side scatter criteria and the GFP positive population determined by applying the appropriate filter using a BD Accuri C6 Plus flow cytometer. The percentage of GFP positive cells was plotted after normalizing to the maximum percentage of GFP positive cells after transduction (measured between 72 - 96 hours after infection).

### Quantitative PCR

Total RNA was recovered from Jurkat cells using an RNAeasy Mini Kit (Qiagen), and cDNA was prepared using an iScript cDNA synthesis kit (BioRad). qPCR for *NOTCH1*, *HES1*, *HES4* and *DTX1* was carried out using a CFX384 Real-Time PCR Detection System. Gene expression was normalized to 28s rRNA as an endogenous control.

### Proteomics Studies: Tandem Affinity Purification

Jurkat cells were transduced with a retrovirus encoding NRARP preceded by an N-terminal tandem HA-FLAG tag and an internal ribosomal entry site (IRES) followed by the coding sequence for the interleukin-2 receptor (IL2R) (pOZ-FH-N) (Nakatani and Ogryzko, 2003). Stable cell lines were generated by magnetic sorting for IL2R-positive cells. For proteomic studies, the resulting cell line was grown in RPMI with 10% bovine growth serum to a density of 3×10^6^ cells/ml and harvested by centrifugation. Cells were lysed with a Dounce homogenizer, and the resulting lysate was immunoprecipitated using anti-FLAG-conjugated agarose beads (Sigma) in a 50 mM Tris buffer, pH 7.5, containing 150 mM NaCl, 1 mM EDTA, 0.5% NP40, 2 mM TCEP and 10% glycerol, supplemented with EDTA-free protease inhibitor tablets (Roche). The beads were washed three times and the immunoprecipitated protein was eluted with a 1xFlag peptide. The eluate was then re-immunoprecipitated with anti-HA-conjugated agarose beads and eluted from the beads with HA peptide for mass spectrometry analysis.

### Mass spectrometry

Protein complexes isolated by tandem affinity purification (Adelmant et al., 2019) were directly processed in solution: Cysteine residues were first reduced with 10 mM dithiothreitol for 30 minutes at 56°C in the presence of 0.1% RapiGest SF (Waters, Milford, MA) and then alkylated with 22.5 mM iodoacetamide for 20 minutes at room temperature in the dark. Proteins were digested overnight at 37°C using 2.5 micrograms of trypsin after adjusting the pH to 8.0 with Tris.

RapiGest SF was cleaved for 30 minutes at 37°C and its by-products were removed by centrifugation. Tryptic peptides were desalted by batch-mode reverse phase solid phase extraction (Poros 10R2) and concentrated in a vacuum concentrator. Peptides were solubilized in 25% acetonitrile containing 0.1% formic acid and further purified by strong cation exchange (Poros 10HS). Peptides were eluted with 25% acetonitrile containing 0.1% formic acid and 300 mM potassium chloride. Acetonitrile was removed using a vacuum concentrator and peptides were reconstituted with 20 μL of 0.1% TFA.

Peptides were loaded onto a precolumn (4 cm POROS 10R2, Applied Biosystems) and eluted with an HPLC gradient (NanoAcquity UPLC system, Waters; 5%–40% B in 90 min; A = 0.2 M acetic acid in water, B = 0.2 M acetic acid in acetonitrile). Peptides were resolved on a self-packed analytical column (50 cm Monitor C18, Column Engineering) and introduced in the mass spectrometer (QExactive HF mass spectrometer, Thermo, Waltham, MA) equipped with a Digital PicoView electrospray source platform (New Objective, Woburn, MA) (Ficarro et al., 2009).

The mass spectrometer was operated in data dependent mode where the top 10 most abundant ions in each MS scan were subjected to high energy collision induced dissociation (HCD, 30% normalized collision energy) and subjected to MS2 scans (isolation width = 1.6 Da, intensity threshold = 2e^5^). Dynamic exclusion was enabled with an exclusion duration of 15 seconds. ESI voltage was set to 3.8 kV.

MS spectra were recalibrated using the background ion (Si(CH_3_)_2_O)_6_ at m/z 445.12 +/− 0.03 and converted into a Mascot generic file format (.mgf) using multiplierz scripts (PMID: 19333238; PMID: 19874609). Spectra were searched using Mascot (version 2.6) against three appended databases consisting of: i) human protein sequences (downloaded from RefSeq on 11/19/2010); ii) common lab contaminants and iii) a decoy database generated by reversing the sequences from these two databases. For Mascot searches, precursor tolerance was set to 15 ppm and product ion tolerance to 25 mmu. Search parameters included trypsin specificity, up to 2 missed cleavages, fixed carbamidomethylation (C, +57 Da) and variable oxidation (M, +16 Da). Spectra matching to peptides from the reverse database were used to calculate a global false discovery rate and were discarded. Data were further processed to remove peptide spectral matches (PSMs) to the forward database with an FDR greater than 1.0%. Peptides shared by two or more genes were excluded from consideration when constructing the final protein list. Any protein identified in more than 1% of 108 negative TAP controls or any of the negative control TAP experiments (PMID: 22810586) was removed from the sets of interactors.

### *In vitro* biotin pull-down assays

Biotin pull-down assays were performed using streptavidin-conjugated agarose (Thermo Fisher) in 20 mM HEPES buffer, pH 7.6, containing 150 mM NaCl, 2 mM TCEP and 0.2% Tween-20. Purified recombinant biotinylated-NRARP, RBPJ and ICN1 (ANK or RAM-ANK) proteins were combined with streptavidin beads at 2 μM and incubated for 30 minutes at room temperature. The beads were washed three times, transferred into gel loading buffer, and the recovered molecules were analyzed by SDS-PAGE using a 4-20% gradient gel followed by staining with Coomassie blue.

### Circular Dichroism Spectroscopy

Circular dichroism (CD) spectra of wild-type, W85E, and W85E/A92W variants of NRARP were acquired at 20 ºC at a protein concentration of 8 μM on a Jasco J-815 instrument in 10 mM phosphate buffer, pH 7.6, containing 150 mM potassium fluoride and 1 mM DTT. Data were acquired in an 0.1 cm pathlength cell and represent the average of 5 scans taken at a 50 nm/min scan rate and a 0.5-nm step size.

### Quantification and Statistical Analysis

Bar graphs display mean ± SD. p values were calculated by one-way ANOVA followed by post hoc Dunnett’s multiple comparison tests where applicable using GraphPad Prism (version 8.0).

### ICN1 stability analysis

Jurkat cells were infected with control retrovirus expressing GFP only, virus expressing dnMAML1, or virus expressing NRARP and grown for 96 hours at 37 C. GFP positive cell populations were then sorted to isolate the GFP positive cell population. Sorted cells were grown for 72 hours, lysed on ice, and probed for total Notch1, activated Notch1 (ICN1), and GAPDH with anti-Notch1 (anti-TAD; Weng et al., 2003), anti-V1744 (Cell Signaling antibody D3B8), and anti-GAPDH (Cell Signaling antibody D16H11) antibodies, respectively.

## Acknowledgments

This work was supported by NIH award R35 CA220340 (to S.C.B.), by a van Maanen Graduate Research Fellowship (to S.M.J.), NIH P01CA203655 (to J.A.M.) and the Dana-Farber Strategic Research Initiative (to J.A.M.). J.C.A. is supported by the Harvard Ludwig Institute and the Michael A. Gimbrone Chair in Pathology at Brigham and Women’s Hospital and Harvard Medical School. The funders had no role in study design, data collection and interpretation, or the decision to submit the work for publication.

