## Supplementary Figures 1 and 2 for "Molecular basis for feedback inhibition of Notch signaling by the Notch regulated ankyrin repeat protein NRARP"

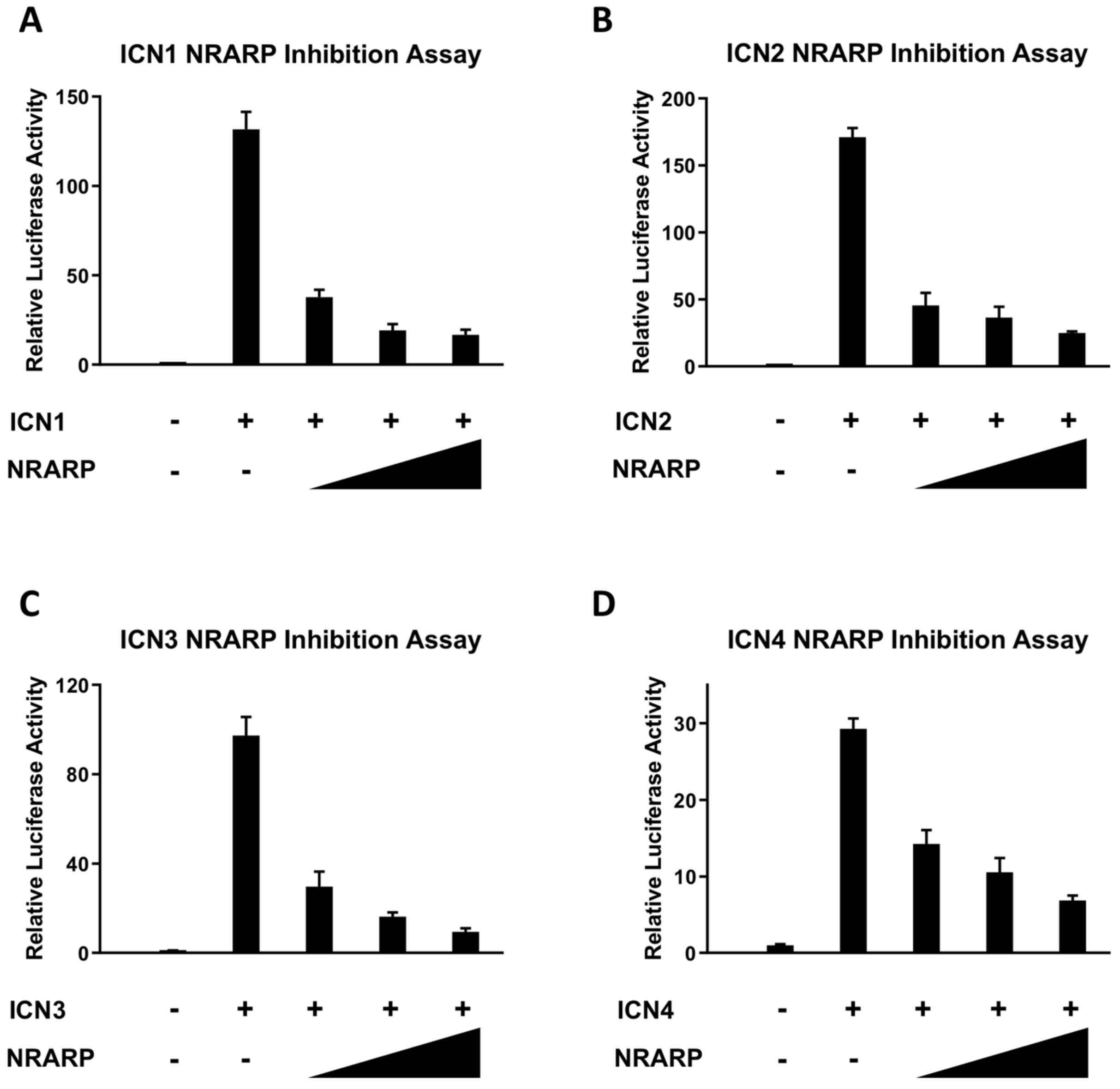


**Supplementary Figure S1.** NRARP inhibits reporter gene induction by all four human Notch receptors. NIH 3T3 cells were transiently transfected with pcDNA3-based plasmids for expression of NRARP and ICN1 (A), ICN2 (B), ICN3 (C), or ICN4 (D), a firefly luciferase reporter plasmid under control of the TP1 Notch-response element, and a plasmid expressing Renilla luciferase. Firefly luciferase activity is reported relative to that of the Renilla luciferase, setting the firefly:Renilla ratio in cells transfected with empty pcDNA vector control to a value of one.

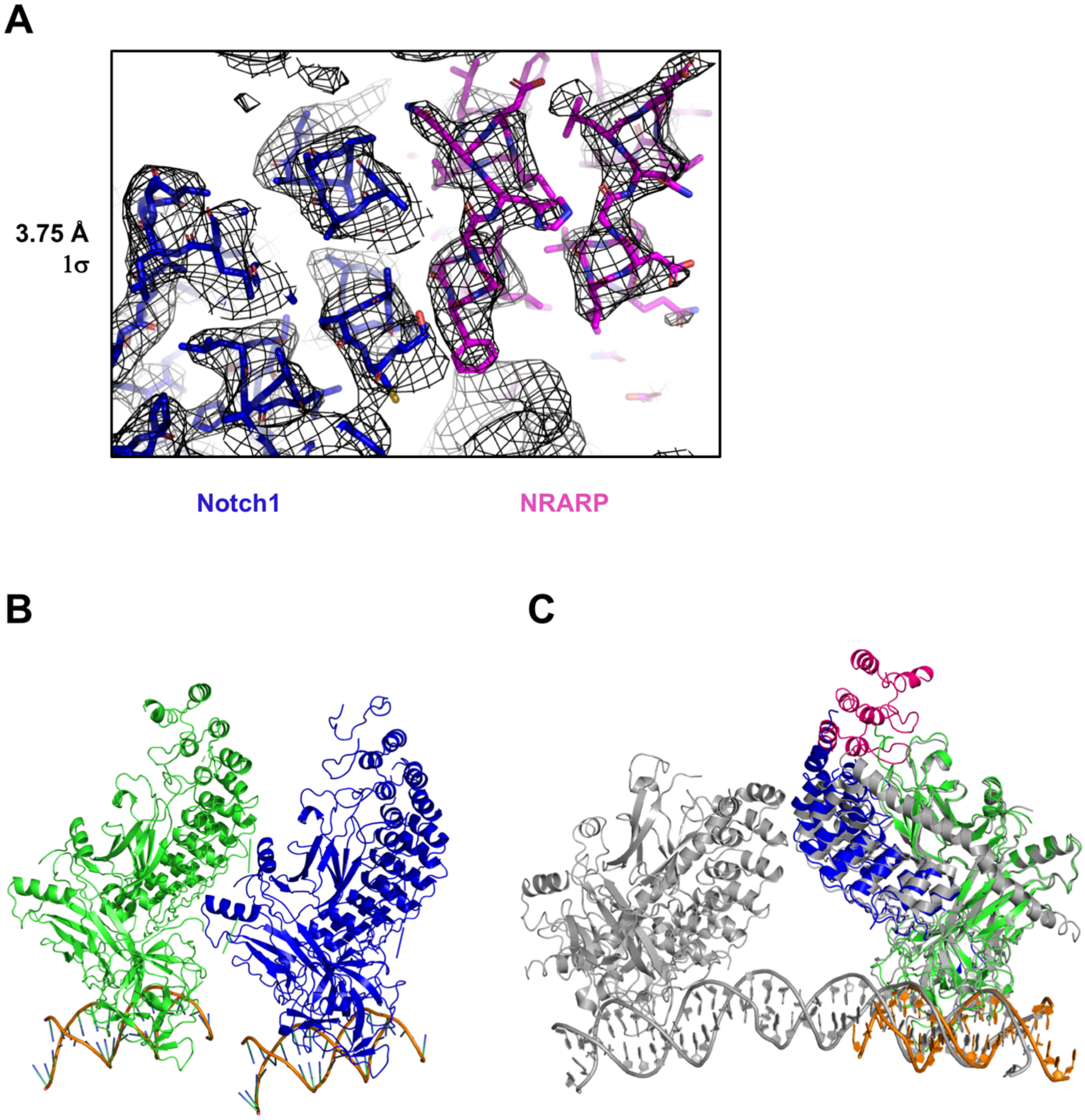


**Supplementary Figure S2.** A. Electron density map of the region at the Notch1-NRARP interface, contoured at 1σ after refinement. B. Ribbon representation of the asymmetric unit of the NRARP-Notch1-RBPJ complex on DNA. The complex contains NRARP (pink), RBPJ (green), Notch1 (blue) and a DNA 16-mer (orange) containing a single RBPJ binding site. **C.** Overlay of the NRARP-NOTCH1-RBPJ complex onto the dimeric RBPJ-Notch1-MAML1-DNA complex (gray; PDB ID code 3NBN).
